# Structure, Dynamics, Receptor Binding, and Antibody Binding of Fully-glycosylated Full-length SARS-CoV-2 Spike Protein in a Viral Membrane

**DOI:** 10.1101/2020.10.18.343715

**Authors:** Yeol Kyo Choi, Yiwei Cao, Martin Frank, Hyeonuk Woo, Sang-Jun Park, Min Sun Yeom, Tristan I. Croll, Chaok Seok, Wonpil Im

## Abstract

The spike (S) protein of severe acute respiratory syndrome coronavirus 2 (SARS-CoV-2) mediates host cell entry by binding to angiotensin-converting enzyme 2 (ACE2), and is considered the major target for drug and vaccine development. We previously built fully-glycosylated full-length SARS-CoV-2 S protein models in a viral membrane including both open and closed conformations of receptor binding domain (RBD) and different templates for the stalk region. In this work, multiple μs-long all-atom molecular dynamics simulations were performed to provide deeper insight into the structure and dynamics of S protein, and glycan functions. Our simulations reveal that the highly flexible stalk is composed of two independent joints and most probable S protein orientations are competent for ACE2 binding. We identify multiple glycans stabilizing the open and/or closed states of RBD, and demonstrate that the exposure of antibody epitopes can be captured by detailed antibody-glycan clash analysis instead of a commonly-used accessible surface area analysis that tends to overestimate the impact of glycan shielding and neglect possible detailed interactions between glycan and antibody. Overall, our observations offer structural and dynamic insight into SARS-CoV-2 S protein and potentialize for guiding the design of effective antiviral therapeutics.

## INTRODUCTION

The outbreak of Coronavirus disease 2019 (COVID-19) caused by severe acute respiratory syndrome coronavirus 2 (SARS-CoV-2) presents a tremendous threat to global public health. It caused over 35 million confirmed cases and more than 1 million deaths as of October, 2020. Due to unavailability of antiviral medicines or approved vaccines, the current treatment strategy is supportive care to relieve symptoms and isolation of infected individuals to reduce transmission, which has placed a huge burden on the public healthcare system and led to massive social and economic distress.

SARS-CoV-2 is an enveloped virus with a positive-sense single-stranded RNA genome^1^. The spike (S) protein anchored in the viral envelope is a class I fusion protein that mediates receptor binding and host cell entry by interacting with human angiotensin converting enzyme-2 (ACE2)^2–4^, and it is also the target of a variety of neutralizing antibodies^5–8^. S protein is a homo-trimer and each monomer has two subunits (S1 and S2) separated by a cleavage site that is recognized by host proteases^9^. A number of recently published structural studies using cryogenic electron microscopy (cryo-EM) have provided a good understanding of the S protein structure at near-atomic resolution^4, 10–12^. The S1 subunit responsible for receptor binding is composed of the signal peptide (SP), N terminal domain (NTD), and receptor binding domain (RBD), and the S2 subunit responsible for membrane fusion is composed of the fusion peptide (FP), two heptad repeats (HR1 and HR2), transmembrane domain (TM), and cytoplasmic domain (CP). The three RBDs on the top of S protein head are conformationally variable. In closed conformations, all three RBDs lay flat with the receptor binding motif occluded by the RBDs on the neighboring monomers. In open conformations, one or multiple RBDs lift up and expose the receptor binding motif(s).

Although the cryo-EM structures of S protein have provided crucial information about its overall structure, highly flexible protein regions such as loops and stalk still remain unresolved. Molecular dynamics (MD) simulation provides molecular-level insight into the underlying mechanisms of biological functions that are difficult to elucidate only with experiments. Recently, cryo-electron tomography (cryo-ET) and MD simulation have been used to explore the conformational variability of S protein stalk that gives the head orientational freedom and allows S protein to scan the host cell surface^13, 14^. However, it still remains unclear what structural freedoms in the stalk portion are determinant to the overall shape of S protein and its orientation, and how they affect the binding to ACE2. In addition, MD simulation along with accessible surface area (ASA) calculations have been used to estimate the impact of glycan shielding on the exposure of antibody epitopes^15^. Mutations of two glycosylation sites have been performed to study the role of two N-linked glycans in stabilizing an RBD open conformation^16^. Further investigation is still required to evaluate whether the ASA difference between glycosylated and non-glycosylated structures truly reflect the impact of glycan shielding on antibody binding, and whether glycans have more functional roles than stabilizing the open-state RBD.

In this work, we present all-atom MD simulations of fully-glycosylated full-length S protein in a viral bilayer, and multiple μs-long trajectories were generated for RBD in open and closed states and S stalk built from different models. We also performed multiple μs-long simulations of non-glycosylated S head-only systems. Our results provide deeper insight into functional roles of glycans that provide not only shielding for immune evasion, but also contribute to the trimer stability and transition of RBD open and close states. Moreover, our simulations give insight into essential structural roles of highly flexible stalk conformations in S protein binding to ACE2.

## RESULTS AND DISCUSSION

### Model structures of fully-glycosylated full-length SARS-CoV-2 S protein

An illustrative snapshot of a fully-glycosylated full-length S protein structure in a viral membrane is shown in Figure 1. When there are many missing residues and domains, the initial models for MD simulations need to be carefully built and validated against available experimental data. We have built the models using GALAXY protein modeling suite^17–19^ for missing residues and domains, ISOLDE^20^ for refining initial models against experimental density maps, and CHARMM-GUI *Glycan Modeler*^21–23^ and *Membrane Builder*^24–26^ for glycan and membrane building. The head of S trimer was built based on cryo-EM structures (PDB ids: 6VSB^4^ and 6VXX^12^). All three chains of 6VXX have RBD in a closed conformation. One chain of 6VSB (A chain) has RBD in an open conformation and the other two chains have RBD in a closed conformation. Two models were selected for each of HR2 linker, HR2-TM, and CP, resulting in a total of 16 structures after the domain by domain assembly. The glycan sequences selected for 22 N-linked and 1 O-linked glycosylation sites of each monomer were based on the mass spectrometry data^27, 28^. The detailed model generation is described in reference^29^. The model name follows the model numbers used for HR2 linker, HR2-TM, and CP structures. For example, “6VSB_1_2_1” represents a model based on 6VSB with HR2 linker model 1 (M1), HR2-TM model 2 (M2), and CP model 1 (M1). All 16 S protein simulation systems and trajectories are available in CHARMM-GUI COVID-19 archive (http://www.charmm-gui.org/docs/archive/covid19).

**Figure 1.**
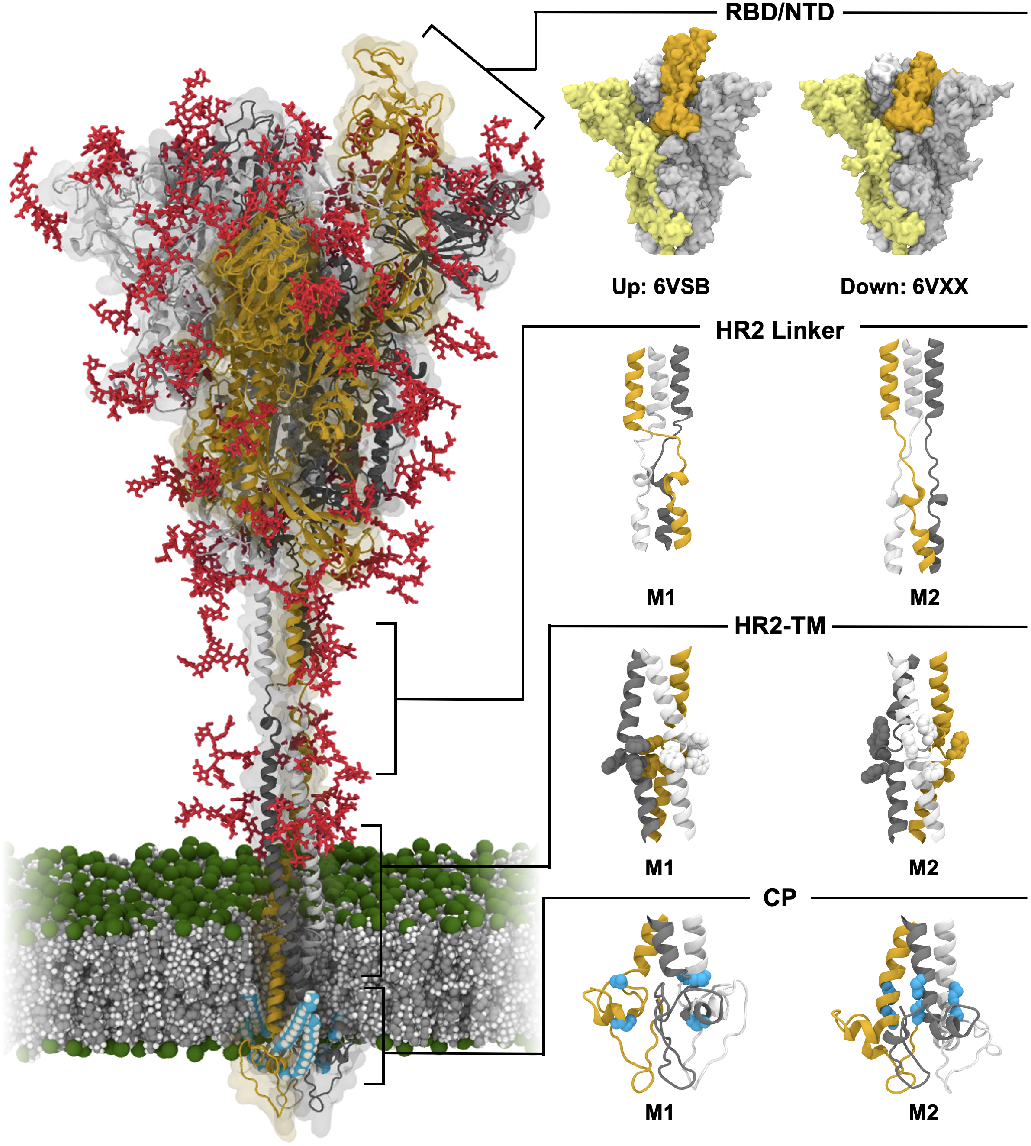
Model structure of fully-glycosylated full-length SARS-CoV-2 S protein in a viral membrane. A model structure of SARS-CoV-2 S protein is shown on the left panel. Two models for RBD/NTD, HR2 linker, HR2-TM, and CP are enlarged on the right panel. The three individual chains of S protein are colored in yellow, gray, and white, respectively, while glycans are represented as red sticks. The palmitoylation sites of S protein are highlighted in cyan. The phosphate, carbon, and hydrogen atoms of the viral membrane are colored in green, gray, and white, respectively. For clarity, water molecules and ions are omitted. All illustrations were created using Visual Molecular Dynamics (VMD)^30^.

### Stalk of S protein consists of two highly flexible linkers

We have performed 1.25-μs all-atom MD simulation of each of 16 models (i.e., a total of 20 μs) each containing about 2.3 million atoms (see Methods). Conformational Analysis Tools (CAT, http://www.md-simulations.de/CAT/) were used for high-throughput analysis of all simulation trajectories. The root-mean-square-deviation (RMSD) time series in **Figure S1** show that the head region of S protein (residue 1-1140) remains stable during the simulations with RMSD of 4 to 5 Å in most systems. The stalk region, however, exhibits highly flexible motions at the HR2 linker and HR2-TM (see Movies S1-2), which is consistent with S protein structures observed in cryo-ET^13^. To further understand the flexible stalk motion, bending characteristics of the two linker regions, defined as θ1 and θ2 (Figure 2A), were quantified. Figure 2B shows the distributions of θ1 and θ2 for each model. Both M1 and M2 of θ1 show similar angle distributions centered at 150° (±15°) and 155° (±12°), respectively. The HR2-TM region, however, exhibited different bending motions. The M1 of HR2-TM shows a narrow distribution centered at 172° (±4°), whereas the M2 of HR2-TM shows a wide distribution centered at 155° (±14°). Twisting motions were also dependent on the HR2-TM model (**Figure S5A**). While both M1 and M2 of HR2-linker show similar twist angle distributions *(φ)* centered at 66° (±46°) and 68° (±49°), respectively, the M1 of HR2-TM shows a narrow distribution centered at 99° (±18°) and the M2 of HR2-TM shows a wide distribution centered at 98° (±71°).

**Figure 2.**
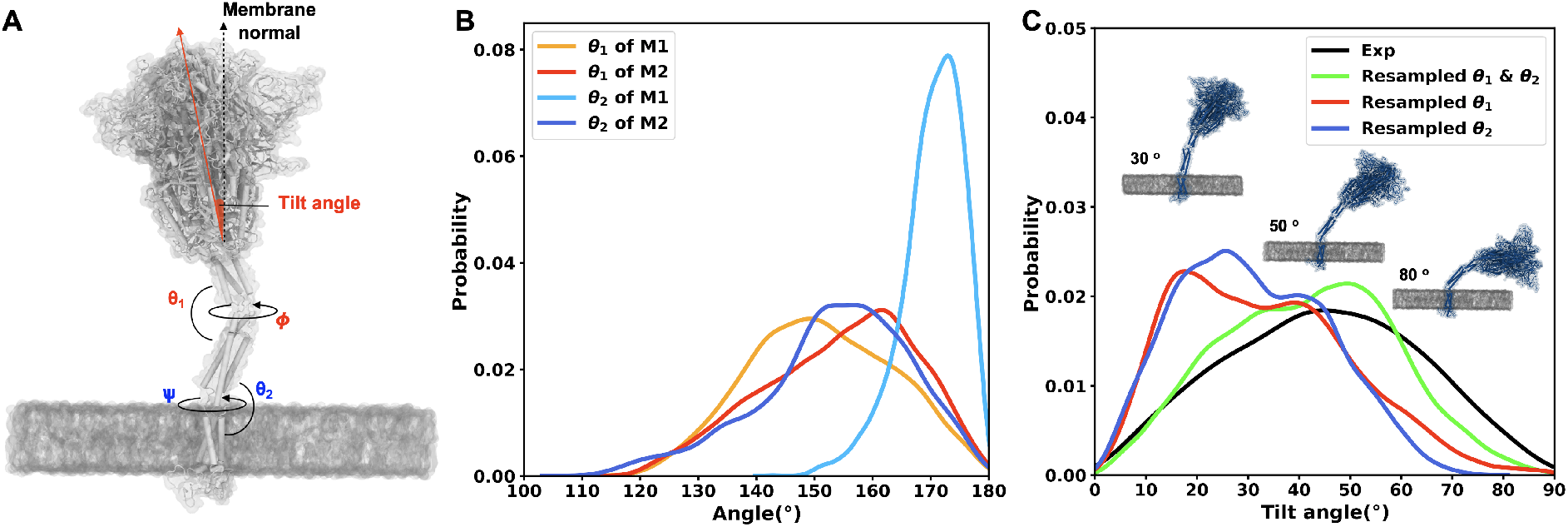
Bending motions of S protein in a viral membrane. (A) Illustrative snapshot of S protein and definition of angles/dihedrals measured to characterize the stalk motion. (B) Probability distribution of bending angle for each HR2 linker and HR2-TM linker model. (C) Probability distributions of tilt angles for the resampled S protein structures compared to the experimental observation^13^. The tilting angle is defined by the principal axis of S protein head and the membrane normal.

These bending and twisting characteristics are consistent with the secondary structure analysis. The secondary structures of HR2 linker M1 and M2 models are mostly in coil conformations during the simulation although local folding and unfolding occur in both models (**Figure S2**). The secondary structure of initial HR2-TM M1 model mainly consists of helical structures that are mostly retained during the simulation time. On the other hand, the secondary structure of M2 initially modeled with turn and bend shows low helicity in the range of L1200-K1215 (**Figure S3**). This indicates that the flexible motions of HR2-TM linker are strongly influenced by the secondary structure and initial model (within the current simulation time). Although the secondary structures of CP domains are different in between two models, they have no significant effect on the motions of stalk (**Figure S4)**. To further characterize the bending motions, the Pearson correlation coefficient (r) was calculated for all combinations of HR2 linker and HR2-TM models (**Figure S5B**). The r values of all cases range from −0.16 to 0.13, indicating that there is no correlation between bending motions of HR2 linker and HR2- TM and thus each linker acts as an independent hinge.

Although 16 x 1.25-μs MD simulations were performed, it does not cover all possible configurations of S protein especially with such flexible two linkers. To increase sampling, utilizing the independent θ1 and θ2 characteristics, S protein orientation was resampled based on three regions: head-HR1, HR1-HR2, and HR2-TM. First, 30 HR2-TM conformations were randomly extracted from each trajectory (excluding HR2-TM M1 models), and their TM domain was superimposed to the TM domain of the initial model to resample HR2 domain motion. Second, 30 HR1-HR2 conformations were randomly extracted and superimposed to each of the resampled HR2 domains. Finally, 30 head-HR1 conformations were randomly extracted and superimposed to the resampled HR1 domains. In summary, 27,000 configurations of S protein on a viral membrane were generated. Figure 2C shows the tilt angle of resampled configurations of S protein using M1 and M2 for HR2 linker and only M2 for HR-TM. The S protein can tilt by up to 90° towards the membrane and tilt angles around 48° are most probable. This tilt angle distribution agrees well with the experimental observation^11, 13^. However, if M1 was used for HR2-TM, the tilt angle distribution of S protein becomes narrow (**Figure S6**). This indicates that both M1 and M2 of HR2 linker are reliable models, but for the HR2-TM, M2 is more appropriate to represent S protein configurations. To further understand the contribution of each independent hinge motion on the tilt angle, S protein was resampled separately with HR2-TM only and with HR1-HR2 only. In both cases, the resampled S protein shows a narrow angle distribution compared to the experimental observation (Figure 2C), indicating that both linkers are necessary for full tilting motions of S protein observed in experiment.

### Flexibility of stalk may facilitate S protein binding to ACE2

To explore the effect of flexible stalk motion on ACE2 binding, we performed structural alignment of S protein to ACE2. The RBD in complex with full-length human ACE2 in the presence of neutral amino acid transporter B^0^AT1 (PDB: 6M17^10^) was used for alignment. Fully independent bending and twisting motions of two stalk linkers allow us to increase the number of S protein samples. 125 head-HR1, HR1-HR2, and HR2-TM-CP conformations were separately extracted from each trajectory with 10-ns interval. Each RBD of head-HR1 conformations was first superimposed to the RBD-ACE2-B^0^AT1 complex. Then, the HR1-HR2 conformations were superimposed to each of HR1 from the previous step. Finally, the HR2-TM-CP conformations were superimposed to each of HR2 from the previous step. Figure 3A shows one of the most probable configurations of S protein-ACE2 complex. The tilting angle *(θ)* is defined in Figure 2, and the distance (*d*) is defined by an arc length between the centers of mass (COMs) of two TM domains. As shown in Figure 3B, *d* ranges from 240 Å to 350 Å and *θ* ranges from 30° to 60°. At the most probable configuration, *d* and *θ* are about 290 Å and 46°, respectively. Note that there is approximately one S protein per 1,000 nm^2^ (316 Å× 316 Å) on the viral surface^14^. This sparse distribution of S protein suggests that receptor binding can be promoted by enough space to have orientational degrees of freedom for RBD. Moreover, it is reported that the most probable tilting angle of prefusion state is about 40° ~ 50°^11, 13^ (also see Figure 2). This tilting angle appears to maximize the accessibility of the receptor binding motif to ACE2 (when the RBD is in an open conformation), which could account for the high infection rate of SARS-CoV-2.

**Figure 3.**
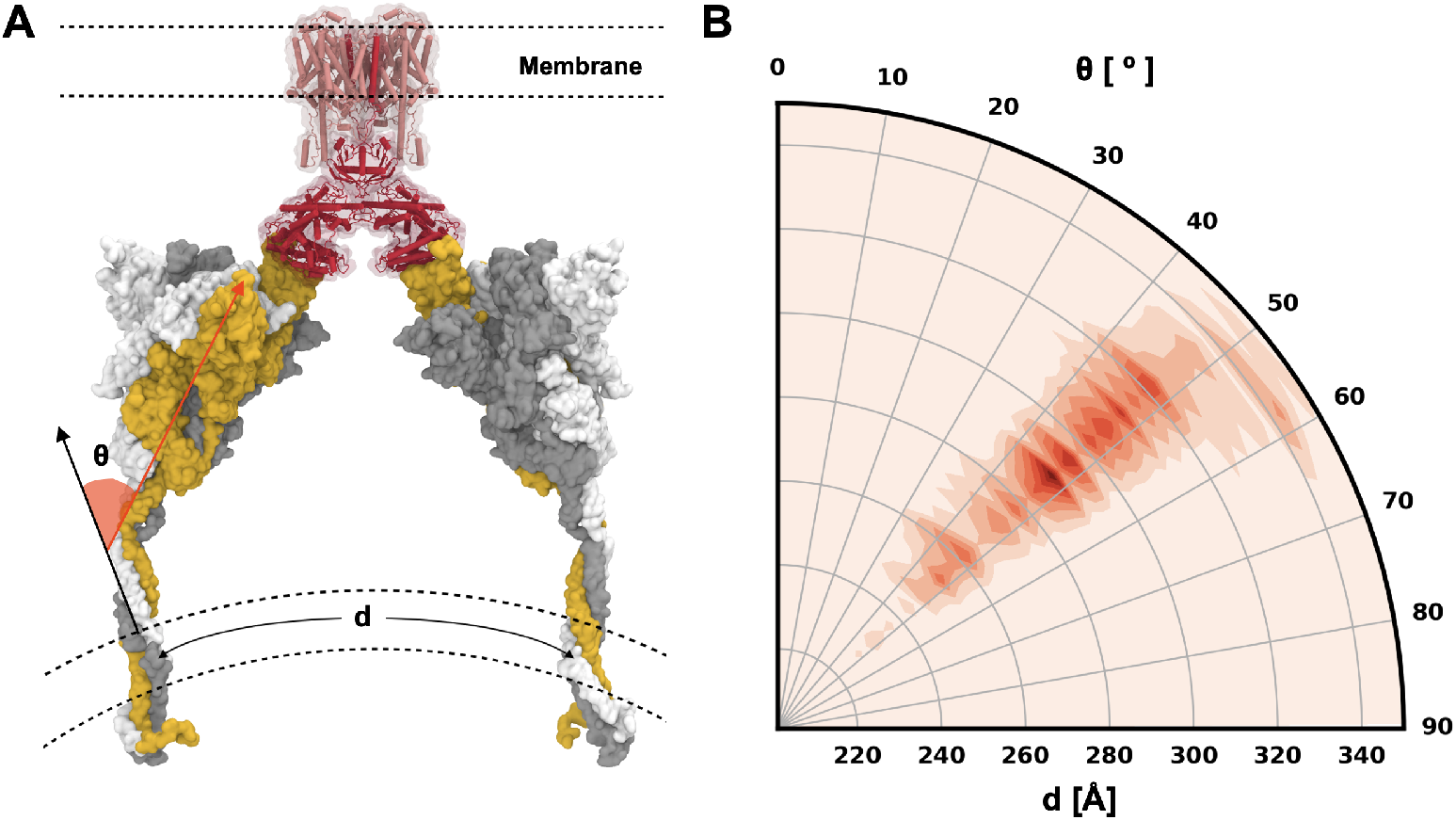
S protein configurations competent for ACE2 binding. (A) Illustrative snapshot of S protein-ACE2-B^0^AT1 complex. Three individual chains of S protein are colored in yellow, gray, and white, and ACE2 and B^0^AT1 are represented as red and pink, respectively. (B) Distribution of tilting angle (*θ*) as a function of the arc length (*d*) between the centers of mass (COMs) of TM domains.

### Glycans influence RBD and NTD motions and contribute to S trimer stability

To explore RBD and NTD motions, we measured two structural features (Figure 4A): RBD-NTD distance (*d*) defined by the minimum distance between RBD and NTD, and RBD orientation angle (*θ*) defined by two points at each end of RBD and the third point on the center axis of S trimer. One RBD (in both open and closed states) forms a U-shaped pocket with the NTD in the same monomer, which is occupied by the neighboring RBD in a closed state. Therefore, *d* estimates the RBD-NTD pocket size and *θ* estimates the extent of RBD opening. The time series of *d* and *θ* in 6VSB and 6VXX are shown in Figures 4B and **S7**, respectively. In cryo-EM structures available in the PDB, *θ* in the open-state RBD ranges from 134° to 153°, and *θ* in the close-state RBD ranges from 88° to 93°. The trajectories of fully-glycosylated models completely cover the RBD orientation angles observed in cryo-EM structures, and explore a wider range of conformational space. In particular, *θ* of 6VSB_A ranges from 120° to 170°, indicating that the open-state RBD is much more flexible than the closed-state RBD, and it is consistent with the RMSD and root-mean-square-fluctuation (RMSF) results shown in **Figure S1**. For both 6VSB_A and 6VSB_C, the simulations cover the NTD-RBD distances observed in the PDB cryo-EM structures. In 6VSB_B, the pocket formed by NTD and RBD (chain B) is empty due to the opening of neighboring RBD (chain A), and consequently, the NTD moved close to RBD.

**Figure 4.**
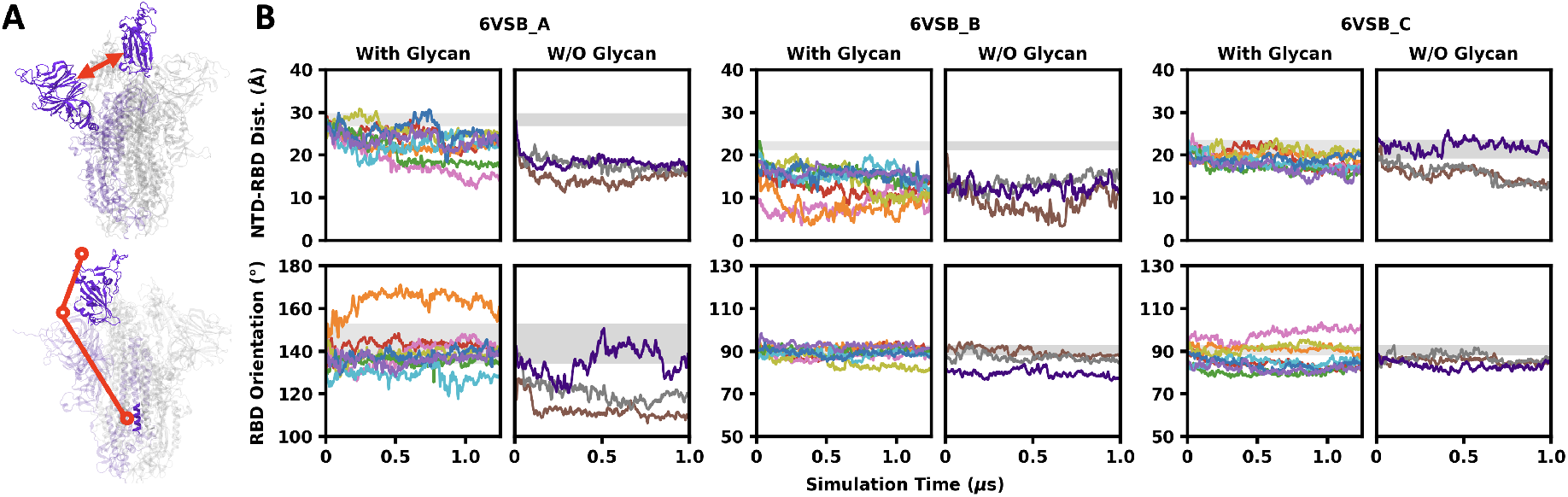
Motions of RBD and NTD in fully-glycosylated systems and non-glycosylated systems. (A) Illustration of NTD-RBD distance *(d)* and RBD orientation angle *(θ).* (B) The time series of *d* and *θ* in three chains of 6VSB. *d* is defined by the minimum distance between RBD (N334 to P527) and NTD (C15 to S305). *θ* is defined by three points corresponding to the (i) COM of L452 and L492, (ii) COM of N334, and (iii) COM of S1030. The ranges of *d* and *θ* observed in available PDB S protein structures are shaded by gray regions.

To investigate the impact of glycans on the transition between RBD open and closed states, we built and simulated non-glycosylated head-only systems (3 replicates for 6VSB and 6VXX) by removing all N-/O-linked glycans and truncating the stalk. It is worth noting that the RBD in non-glycosylated 6VSB_A started to close at the very beginning of trajectories in all three replicas (Figure 4B). In the trajectory (colored in brown), *θ* decreases to 110°, which is about in the middle of open and closed states, and in the trajectory (colored in purple), the RBD reverted to opening at around 0.25 μs. Since the transition between RBD open and closed states is a complicated process involving the motions of multiple domains and attached glycans, it may require much longer simulation time to capture the conformation in which the geometries of both protein and glycan satisfy the requirement for the transition to occur. Nonetheless, dramatic transitions from RBD open to more closed states in non-glycosylated 6VSB_A indicates that the RBD open state is unstable when the glycans are removed. This implies that glycosylation has critical roles in the viral entry as the RBD needs to be open in order to interact with ACE2.

We identified three N-glycans that have important roles in the conformational change of RBD. They are attached to N165 and N234 on NTD and N343 on RBD, respectively. When RBD is open (6VSB_A), N165 and N234 glycans on the NTD of neighboring chain (6VSB_B) are both located below the open-state RBD (Figure 5A), which holds the open-state RBD. Although both are near the open-state RBD, only N165 glycan has frequent contacts with RBD (>85% of snapshots) (Figure 5C), and N234 glycan interacts with the open-state RBD occasionally (<5% of snapshots), which is different from the findings from Casalino et al’s study^16^. Such a difference could attribute to the initial models and simulation lengths. When RBD is closed (all except 6VSB_A), the RBD forms a sandwich-like arrangement with two glycans. N165 glycan is located above the RBD and N234 glycan is located below the RBD (Figure 5B). Both glycans frequently interact with the closed-state RBD (Figure 5C), which makes transition to an open conformation hard. The glycan attached to N343 on RBD orients toward the solvent and hardly interacts with other domains when RBD is open (6VSB_A) (Figure 5D). When an RBD is closed (6VSB_B) but the neighboring RBD (6VSB_A) is open, N343 glycan orients toward the pocket between NTD and RBD and interacts with N165 glycan, which makes open to closed state transition of neighboring RBD difficult. When RBD is closed (all expect 6VSB_A and 6VSB_B) and the neighboring RBD is also closed, N343 glycan makes extensive interactions with the neighboring RBD (Figure 5E), which contributes to the stability of both RBDs in closed conformation The atom contacts between RBD N343 glycan and neighboring RBD exist in more than 95% of snapshots when both RBDs are closed (Figure 5F). Given that RBD is shown to constantly transit between open and closed states in experiment, we propose that glycans serve as a clutch that temporarily holds the RBD in an open or closed conformation, which modulates the lifetime of both open and closed states.

**Figure 5.**
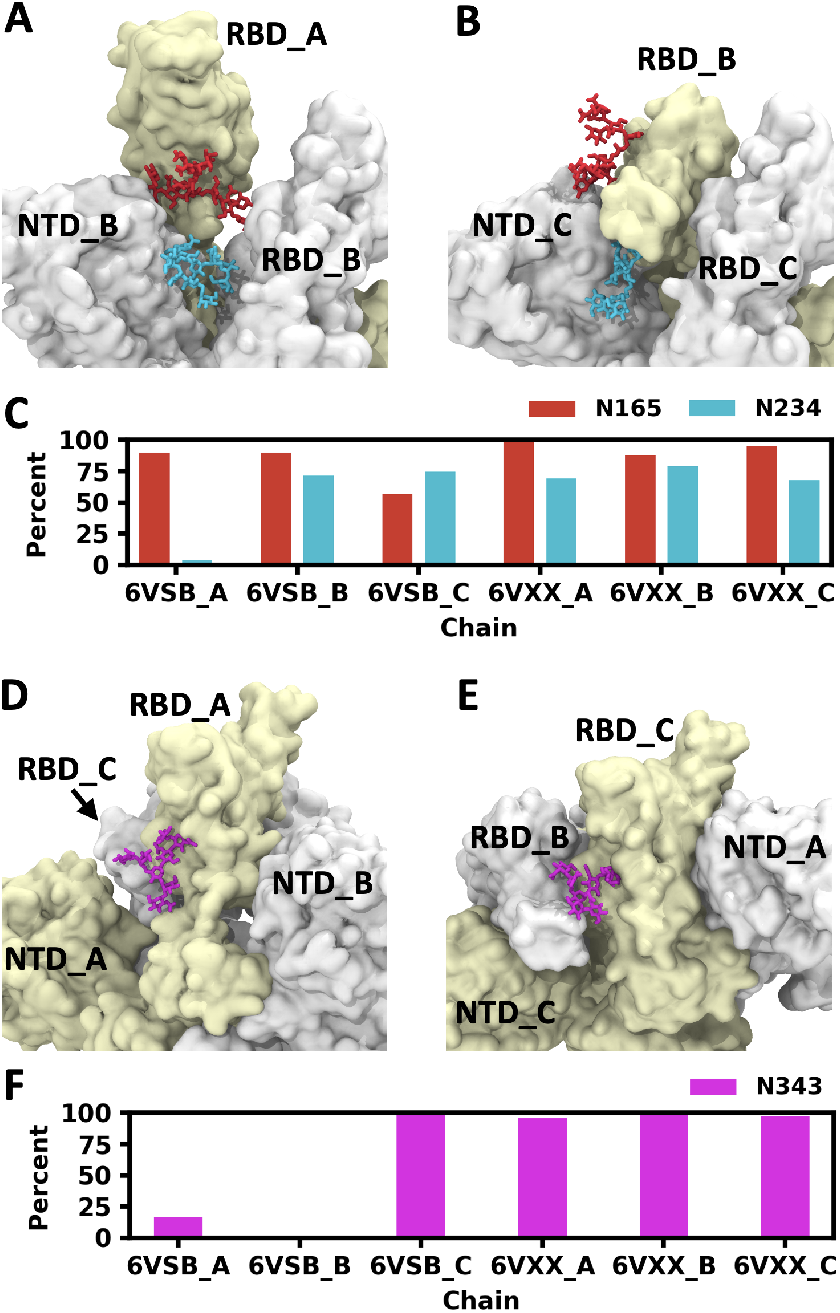
Critical roles of N165, N234, and N343 glycans in the transition between RBD open and closed conformations. (A) The open-state RBD (6VSB_A, pale yellow) is above the N165 (red) and N234 (cyan) glycans on the neighboring NTD. (B) The closed-state RBD (6VSB_B) is between N165 and N234 glycans. (C) Frequency of contacts between RBD and two glycans (N165 and N234). (D) N343 (magenta) glycan on the open-state RBD (6VSB_A) is free. (E) N343 glycan on the closed-state RBD (6VSB_C) interacts with the neighboring closed-state RBD (6VSB_C). (F) Frequency of contacts between N343 glycan and the neighboring RBD.

In addition, we calculated the accessible surface area (ASA) reduced due to formation of S trimer. For example, the ASA reduction for chain A was calculated by S_A_ + S_BC_ - S_ABC_, where S_A_, S_BC_, and S_ABC_ are the ASA of chain A, chains BC-only complex, and chains ABC complex (i.e., S trimer), respectively. The ASA reduction due to trimer formation was split into the portion from protein only and the portion involving glycans. The trimer interface interactions involving glycans is about 30% when the entire S1 subunit is considered, and it increases to about 40% when only RBD and NTD are considered (**Figure S8**). This suggests that glycans makes significant contributions to the stability of S trimer, which is different to the common belief that protein-protein interactions are the only dominating factor to the stability of a protein multimer.

### The impact of glycan shielding on antibody binding is overestimated

Viruses evolve to minimize the immunogenicity by coating the exposed viral proteins with non-immunogenic or weakly-immunogenic glycans. It is commonly believed that the glycans on viral envelope shield viruses from host immune system. To get an impression of such glycan shielding, we aligned the S head in each trajectory snapshot and the glycan distributions are shown in Figure 6A. Most glycans are very flexible and they move around in a wide range of space, which covers most of the trimer surface. However, the role of glycan is not limited to shielding. During the past decade, many glycan-dependent HIV neutralizing antibodies have been discovered and extensively studied, which target both envelope protein and glycans^31^’^33^. In the cryo-EM structure of S trimer in complex with S309 antibody (PDB ids: 6WPS and 6WPT^34^), S309 interacts with the glycan attached to N343.

**Figure 6.**
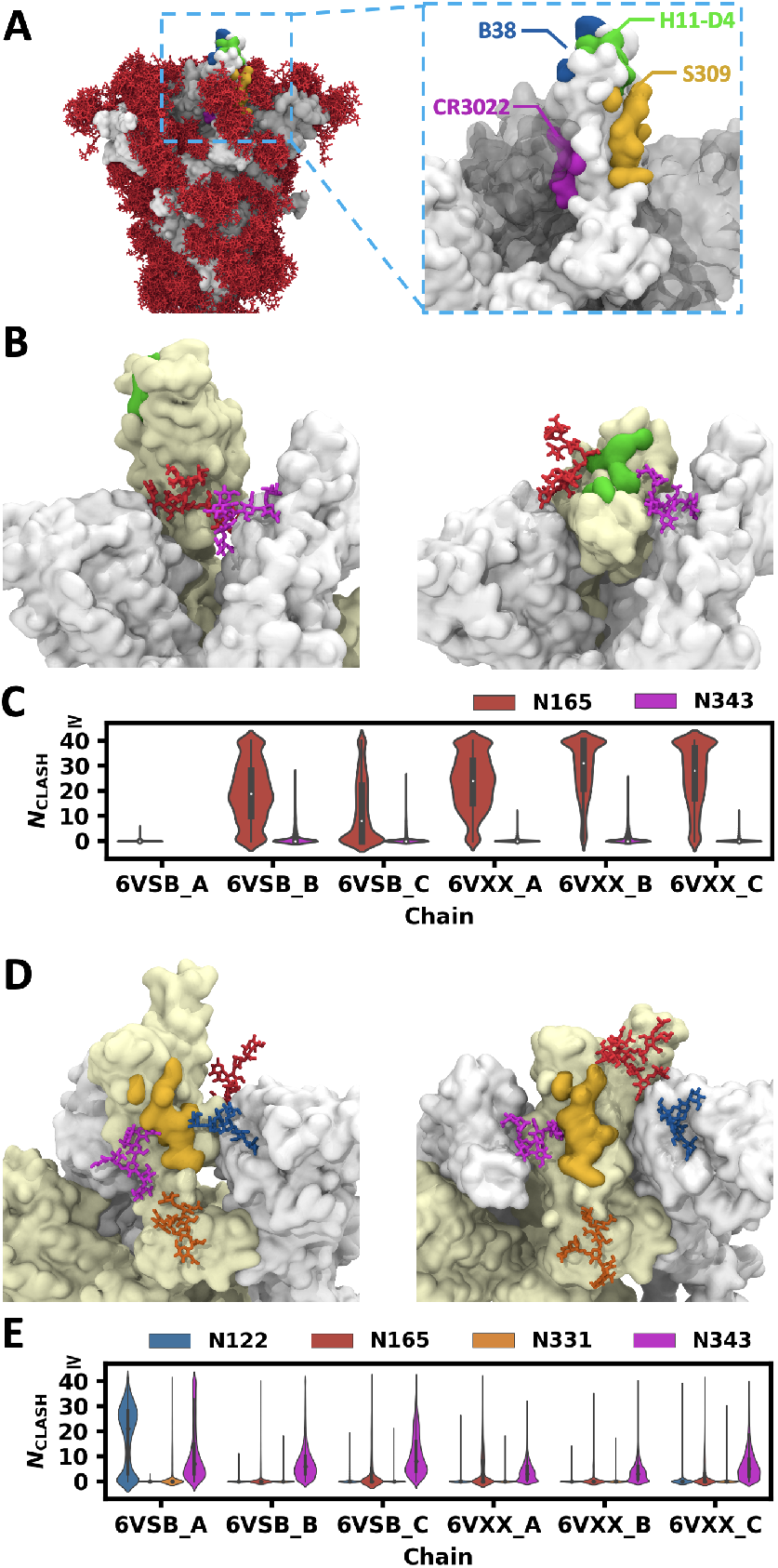
Clash between glycans and superimposed antibodies. (A) Distribution of glycans when S head structures in multiple snapshots are aligned (left). Four epitopes targeted by neutralizing antibodies are shown in different colors (right). (B) H11-D4 epitope (green) in RBD (left: open, right: closed), N165 glycan (red) on the neighboring NTD, and N343 glycan (violet) on the neighboring RBD. (C) Distributions of glycan heavy atom numbers in clash (*N*_CLASH_) with the superposed H11-D4 antibody. (D) S309 epitope (yellow), N331 (orange) and N343 (violet) glycans on RBD (left: open, right: closed), N122 (blue) and N165 (red) glycans on the neighboring NTD. (E) Distributions of *N*_CLASH_ with the superimposed S309 antibody.

To explore the role of glycans in antibody binding, we used TM-align^35^ to superimpose the RBD in RBD-antibody complex structures in the PDB onto the RBD in each simulation snapshot, and calculated the number of glycan heavy atoms that have clashes with the antibodies (*N*_CLASH_, **Figure S9**). In this work, we discuss four RBD-targeting antibodies, namely B38, CR3022, H11-D4, and S309 (Figure 6A). The epitopes of B38 and CR3022 are irrelevant to glycans. B38 binds to the same region of RBD as ACE2 does. This epitope is fully exposed in the open-state RBD, but when the RBD is closed, B38 epitope is masked by the neighboring RBD in either open or closed state (**Figure S10A**). The epitope of CR3022 is in the inner surface of RBD, and it is only accessible when all three RBD are open (**Figure S10B**).

The epitope of H11-D4 is next to the epitope of B38, and it is also fully exposed when RBD is open. When RBD is closed, the glycans attached to N165 and N343 on the neighboring chain are located near this epitope (Figures 6B and **S10C**). As shown in the distribution of *N*_CLASH_ with H11-D4 (Figure 6C), N343 glycan rarely interferes with the antibody, but N165 glycan has high probabilities to make severe clashes with the antibody when RBD is closed. For comparison, we also aligned a nanobody to the RBD in each simulation snapshot. Though N165 glycan still frequently makes clashes with the nanobody, the frequency of severe clashes is much lower (**Figure S11**) and thus there is a chance a nanobody can bind to this epitope as shown in PDB id 6Z43.

The epitope of S309 is surrounded by four glycans. Two of them are attached to N331 and N343 on the same RBD, and the other two are attached to N122 and N165 on the neighboring NTD (Figures 6D and **S10D**). The distributions of *N*_CLASH_ with S309 are shown in Figure 6E. N165 and N331 glycans rarely interfere with S309 antibody in both open- and closed-state RBD. N343 glycan has minor clashes with the antibody in most snapshots, and such minor clashes are not sufficient to block antibody binding, as these clashes can be easily removed with small changes in glycan conformation and orientation. The antibody-glycan interactions can also contribute to the antibody binding, which is observed in the cryo-EM structures. In more than half of all snapshots, N122 glycan has severe clashes with the antibody when RBD is open, but it moves away from the superposition of antibody in the remaining snapshots. This suggests that S309 epitope in the open state RBD is blocked by N122 glycan in more than half of the simulation time, but it is still accessible to the antibody.

As a comparison, we calculated the ASA of S309 and H11-D4 epitopes using a probe radius of 7.2 Å that is commonly used to approximate the size of hypervariable loops of antibody, and compared the epitope ASA with and without glycans (**Figure S12**). For H11-D4, we observed significant decreases of the epitope ASA in all chains except 6VSB_A and 6VSB_C, which is generally consistent with the frequency of clashes between glycans and antibodies. However, for S309, the epitope ASA decreases significantly in all chains when glycans are present. This is contradictory to the result that only N343 glycans occasionally have only slight clashes with the superimposed antibody when the RBD is close (all except 6VSB_A). In addition, the PDB structures of S trimer in complex with S309 shows that N343 glycan interacts with the antibody. In the calculation of ASA, a point on the surface is considered inaccessible even if the probe sphere has a very tiny clash with the molecule. However, the shape of antibody is not a sphere and it can have narrow shaped regions that extends deep into the pocket in the epitope. A glycan like the one attached to N343 can reduce the epitope ASA even though it may contribute to antibody binding. Therefore, in some cases, simple comparison of protein ASA with and without glycans is likely to overestimate the impact of glycan shielding.

## CONCLUSIONS

In this work, we present multiple μs-long all-atom MD simulations of fully-glycosylated fulllength S protein in a viral membrane. Our MD simulations reveal the overall shape of S protein and its orientation on the membrane surface are determined by highly flexible stalk composed of two independent joints. Importantly, S protein models from our simulation allow us to predict possible configurations of S protein-ACE2-B^0^AT1 complex with allowable orientations and distances between two S proteins on the membrane surface. The simulation here also provides insight into how glycans influence the open/closed state change of RBD and the antibody binding to RBD epitopes. We identify glycans attached to multiple glycosylation sites that stabilize the open and/or closed states of RBD by making a high energetic barrier between the open-closed transition. The simulation of non-glycosylated systems shows that the open-state RBD becomes unstable when glycans are removed and the transition to close state occurred at the early stage of simulation. By aligning RBD-antibody complex structures to the simulation trajectories, we reveal that the impact of glycan shielding is overestimated by a simple ASA analysis. More importantly, the glycan does not only serve as shielding for immune evasion, but it can also contribute to antibody binding. Our work sheds light on the full structure and dynamics of S protein and we hope our work to be useful for development of vaccines and antiviral drugs.

## METHODS

In this study, the CHARMM36(m) force field was used for proteins^36^, lipids^37, 38^, and carbohydrates^39^’^41^. TIP3P water model^42^ was utilized along with 0.15 M KCl solution. The total number of atoms is 2,343,394 (6VSB_1_1_1: 668,899 water molecules, 2,128 K^+^, and 1,857 Cl’); see CHARMM-GUI COVID-19 Archive (http://www.charmm-gui.org/docs/archive/covid19) for other system information. The van der Waals interactions were smoothly switched off over 10-12 Å by a force-based switching function^43^ and the long-range electrostatic interactions were calculated by the particle-mesh Ewald method^44^ with a mesh size of ~1 Å.

All simulations were performed using the inputs generated by CHARMM-GUI^45^ and GROMACS 2018.6^46^ for both equilibration and production with LINCS algorithm^47^. Temperature was maintained using a Nosé-Hoover temperature coupling method^48, 49^ with a τt of 1 ps, for pressure coupling (1 bar), semi-isotropic Parrinello-Rahman method^50, 51^ with a τp of 5 ps and compressibility of 4.5 x 10’^5^ bar’^1^ was used. During equilibration run, NVT (constant particle number, volume, and temperature) dynamics was first applied with a 1-fs time step for 250 ps. Subsequently, the NPT (constant particle number, pressure, and temperature) ensemble was applied with a 1-fs time step (for 2 ns) and with a 2-fs time step (for 18 ns). During the equilibration, positional and dihedral restraint potentials were applied, and their force constants were gradually reduced. Production run was performed with a 4-fs time-step using the hydrogen mass repartitioning technique^52^ without any restraint potential. Each system ran about 20 ns / day with 1,024 CPU cores on NURION in the Korea Institute of Science and Technology Information.

## ACKNOWLEDGMENT

This study was supported in part by grants from NIH GM126140, GM138472, NSF OAC-1931343, DBI-2011234, DBI-1660380, and MCB-1810695, a Friedrich Wilhelm Bessel Research Award from the Humboldt Foundation (WI), National Research Foundation of Korea grants (2019M3E5D4066898 and 2016M3C4A7952630) funded by the Korea government (CS), Wellcome Trust grant number 209407/Z/17/Z (TC), and the National Supercomputing Center with supercomputing resources including technical support (KSC-2020-CRE-0089 and KSC-2020-CRE-0094) (MSY and CS).

## NOTES

The authors declare no competing financial interest.

## Notes

http://www.charmm-gui.org/docs/archive/covid19

### Competing Interest Statement

The authors have declared no competing interest.

## REFERENCES

1. Lai, M. M. & Cavanagh, D. The molecular biology of coronaviruses. Adv. Virus Res. 48, 1–100 (1997).

2. Letko, M., Marzi, A. & Munster, V. Functional assessment of cell entry and receptor usage for SARS-CoV-2 and other lineage B betacoronaviruses. Nat. Microbiol. 5, 562–569 (2020).

3. Hoffmann, M. et al. SARS-CoV-2 cell entry depends on ACE2 and TMPRSS2 and is blocked by a clinically proven protease inhibitor. Cell (2020).

4. Wrapp, D. et al. Cryo-EM structure of the 2019-nCoV spike in the prefusion conformation. Science 367, 1260–1263 (2020).

5. Cao, Y. et al. Potent neutralizing antibodies against SARS-CoV-2 identified by high-throughput single-cell sequencing of convalescent patients’ B cells. Cell (2020).

6. Yuan, M. et al. A highly conserved cryptic epitope in the receptor binding domains of SARS-CoV-2 and SARS-CoV. Science 368, 630–633 (2020).

7. Shi, R. et al. A human neutralizing antibody targets the receptor binding site of SARS-CoV-2. Nature 584, 120–124 (2020).

8. Ju, B. et al. Human neutralizing antibodies elicited by SARS-CoV-2 infection. Nature 584, 115–119 (2020).

9. Hoffmann, M., Kleine-Weber, H. & Pöhlmann, S. A multibasic cleavage site in the spike protein of SARS-CoV-2 is essential for infection of human lung cells. Mol. Cell (2020).

10. Yan, R., Zhang, Y., Li, Y., Xia, L., Guo, Y. & Zhou, Q. Structural basis for the recognition of SARS-CoV-2 by full-length human ACE2. Science 367, 1444–1448 (2020).

11. Yao, H. et al. Molecular architecture of the SARS-CoV-2 virus. Cell 183, 1–9 (2020).

12. Walls, A. C., Park, Y.-J., Tortorici, M. A., Wall, A., McGuire, A. T. & Veesler, D. Structure, function, and antigenicity of the SARS-CoV-2 spike glycoprotein. Cell 181, 281–292 (2020).

13. Turoňová, B. et al. In situ structural analysis of SARS-CoV-2 spike reveals flexibility mediated by three hinges. Science 370, 203–208 (2020).

14. Ke, Z. et al. Structures and distributions of SARS-CoV-2 spike proteins on intact virions. Nature (2020).

15. Grant, O. C., Montgomery, D., Ito, K. & Woods, R. J. Analysis of the SARS-CoV-2 spike protein glycan shield: implications for immune recognition. bioRxiv (2020).

16. Casalino, L. et al. Beyond Shielding: The Roles of Glycans in the SARS-CoV-2 Spike Protein. ACS Central Science (2020).

17. Ko, J., Park, H. & Seok, C. GalaxyTBM: template-based modeling by building a reliable core and refining unreliable local regions. BMC Bioinf. 13, 198 (2012).

18. Ko, J., Lee, D., Park, H., Coutsias, E. A., Lee, J. & Seok, C. The FALC-Loop web server for protein loop modeling. Nucleic Acids Res. 39, W210–W214 (2011).

19. Park, T., Baek, M., Lee, H. & Seok, C. GalaxyTongDock: Symmetric and asymmetric ab initio protein-protein docking web server with improved energy parameters. J. Comput. Chem. 40, 2413–2417 (2019).

20. Croll, T. I. ISOLDE: a physically realistic environment for model building into low-resolution electron-density maps. Acta Crystallogr., Sect. D: Struct. Biol. 74, 519–530 (2018).

21. Jo, S., Song, K. C., Desaire, H., MacKerell Jr, A. D. & Im, W. Glycan Reader: automated sugar identification and simulation preparation for carbohydrates and glycoproteins. J. Comput. Chem. 32, 3135–3141 (2011).

22. Park, S.-J. et al. Glycan Reader is improved to recognize most sugar types and chemical modifications in the Protein Data Bank. Bioinformatics 33, 3051–3057 (2017).

23. Park, S.-J. et al. CHARMM-GUI Glycan Modeler for modeling and simulation of carbohydrates and glycoconjugates. Glycobiology 29, 320–331 (2019).

24. Jo, S., Kim, T. & Im, W. Automated builder and database of protein/membrane complexes for molecular dynamics simulations. PloS one 2, e880 (2007).

25. Jo, S., Lim, J. B., Klauda, J. B. & Im, W. CHARMM-GUI Membrane Builder for mixed bilayers and its application to yeast membranes. Biophys. J. 97, 50–58 (2009).

26. Wu, E. L. et al. CHARMM-GUI membrane builder toward realistic biological membrane simulations. J. Comput. Chem. 35, 1997–2004 (2014).

27. Watanabe, Y., Allen, J. D., Wrapp, D., McLellan, J. S. & Crispin, M. Site-specific glycan analysis of the SARS-CoV-2 spike. Science (2020).

28. Shajahan, A., Supekar, N. T., Gleinich, A. S. & Azadi, P. Deducing the N-and O-glycosylation profile of the spike protein of novel coronavirus SARS-CoV-2. bioRxiv (2020).

29. Woo, H. et al. Modeling and Simulation of a Fully-glycosylated Full-length SARS-CoV-2 Spike Protein in a Viral Membrane. J. Phys. Chem. B 124, 7128–7137 (2020).

30. Humphrey, W., Dalke, A. & Schulten, K. VMD: visual molecular dynamics. J. Mol. Graphics 14, 33–38 (1996).

31. McLellan, J. S. et al. Structure of HIV-1 gp120 V1/V2 domain with broadly neutralizing antibody PG9. Nature 480, 336–343 (2011).

32. Walker, L. M. et al. Broad neutralization coverage of HIV by multiple highly potent antibodies. Nature 477, 466–470 (2011).

33. Walker, L. M. et al. Broad and potent neutralizing antibodies from an African donor reveal a new HIV-1 vaccine target. Science 326, 285–289 (2009).

34. Pinto, D. et al. Structural and functional analysis of a potent sarbecovirus neutralizing antibody. BioRxiv (2020).

35. Zhang, Y. & Skolnick, J. TM-align: a protein structure alignment algorithm based on the TM-score. Nucleic Acids Res. 33, 2302–2309 (2005).

36. Huang, J. et al. CHARMM36m: an improved force field for folded and intrinsically disordered proteins. Nat. Methods 14, 71–73 (2017).

37. Klauda, J. B. et al. Update of the CHARMM all-atom additive force field for lipids: validation on six lipid types. J. Phys. Chem. B 114, 7830–7843 (2010).

38. Klauda, J. B., Monje, V., Kim, T. & Im, W. Improving the CHARMM force field for polyunsaturated fatty acid chains. J. Phys. Chem. B 116, 9424–9431 (2012).

39. Guvench, O. et al. Additive empirical force field for hexopyranose monosaccharides. J. Comput. Chem. 29, 2543–2564 (2008).

40. Guvench, O., Hatcher, E., Venable, R. M., Pastor, R. W. & MacKerell Jr, A. D. CHARMM additive all-atom force field for glycosidic linkages between hexopyranoses. J. Chem. Theory Comput. 5, 2353–2370 (2009).

41. Hatcher, E., Guvench, O. & MacKerell Jr, A. D. CHARMM additive all-atom force field for aldopentofuranoses, methyl-aldopentofuranosides, and fructofuranose. J. Phys. Chem. B 113, 12466–12476 (2009).

42. Jorgensen, W. L., Chandrasekhar, J., Madura, J. D., Impey, R. W. & Klein, M. L. Comparison of simple potential functions for simulating liquid water. J. Chem. Phys. 79, 926–935 (1983).

43. Steinbach, P. J. & Brooks, B. R. New spherical-cutoff methods for long-range forces in macromolecular simulation. J. Comput. Chem. 15, 667–683 (1994).

44. Essmann, U., Perera, L., Berkowitz, M. L., Darden, T., Lee, H. & Pedersen, L. G. A smooth particle mesh Ewald method. J. Chem. Phys. 103, 8577–8593 (1995).

45. Lee, J. et al. CHARMM-GUI input generator for NAMD, GROMACS, AMBER, OpenMM, and CHARMM/OpenMM simulations using the CHARMM36 additive force field. J. Chem. Theory Comput. 12, 405–413 (2016).

46. Van Der Spoel, D., Lindahl, E., Hess, B., Groenhof, G., Mark, A. E. & Berendsen, H. J. GROMACS: fast, flexible, and free. J. Comput. Chem. 26, 1701–1718 (2005).

47. Hess, B., Bekker, H., Berendsen, H. J. & Fraaije, J. G. LINCS: a linear constraint solver for molecular simulations. J. Comput. Chem. 18, 1463–1472 (1997).

48. Nosé, S. A molecular dynamics method for simulations in the canonical ensemble. Mol. Phys. 52, 255–268 (1984).

49. Hoover, W. G. Canonical dynamics: Equilibrium phase-space distributions. Phys. Rev. A 31, 1695 (1985).

50. Parrinello, M. & Rahman, A. Polymorphic transitions in single crystals: A new molecular dynamics method. J. Appl. Phys. 52, 7182–7190 (1981).

51. Nosé, S. & Klein, M. Constant pressure molecular dynamics for molecular systems. Mol. Phys. 50, 1055–1076 (1983).

52. Hopkins, C. W., Le Grand, S., Walker, R. C. & Roitberg, A. E. Long-time-step molecular dynamics through hydrogen mass repartitioning. J. Chem. Theory Comput. 11, 1864–1874 (2015).

