## Supporting Figures for "Structure, Dynamics, Receptor Binding, and Antibody Binding of Fully-glycosylated Full-length SARS-CoV-2 Spike Protein in a Viral Membrane"

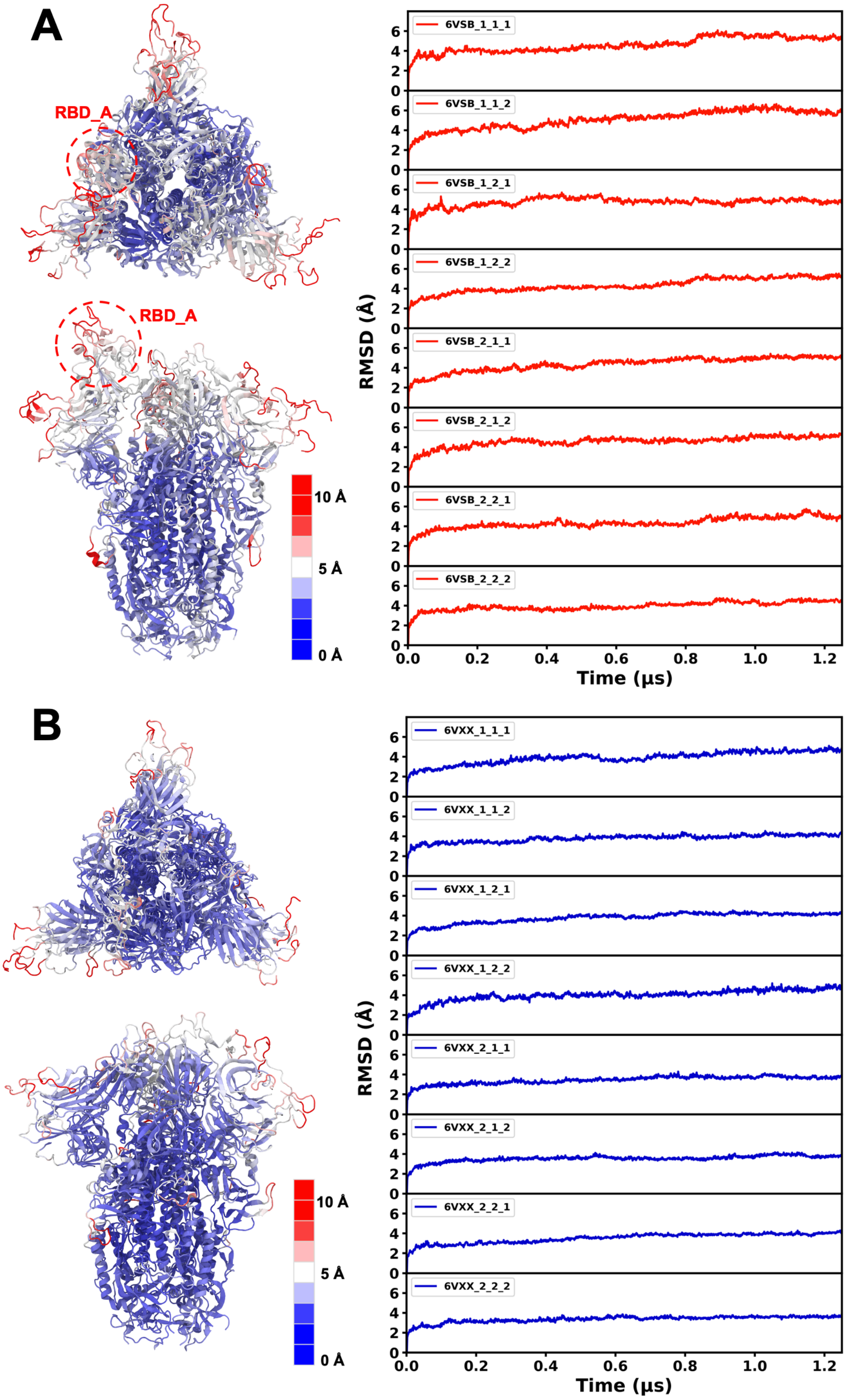


**Figure S1. RMSF and RMSD of S protein head.** (left) Root-mean-square-fluctuation (RMSF) and (right) time series of root-mean-square-deviation (RMSD) of S protein head (residue 1-1140) for (A) RDB up and (B) down conformations of chain A (RBD_A).

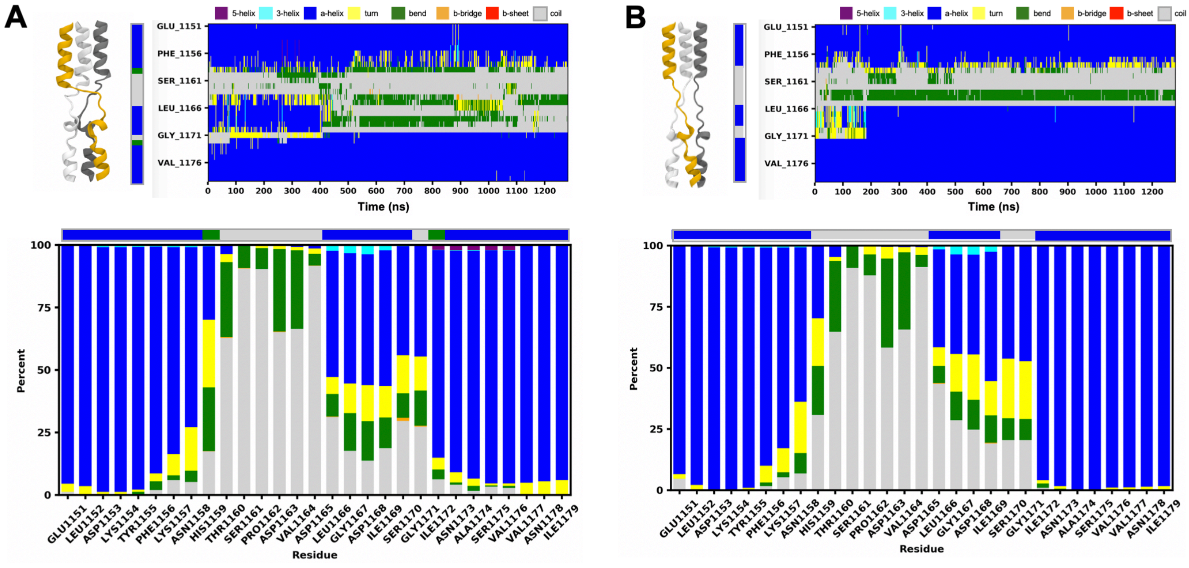


**Figure S2. Secondary structure analysis of HR2 linker (A) model 1 and (B) model 2.** (Top from right) a snapshot of model structure, an initial secondary structure, and time series of secondary structures of a representative system, and (bottom) secondary structure contents per residue from all simulations.

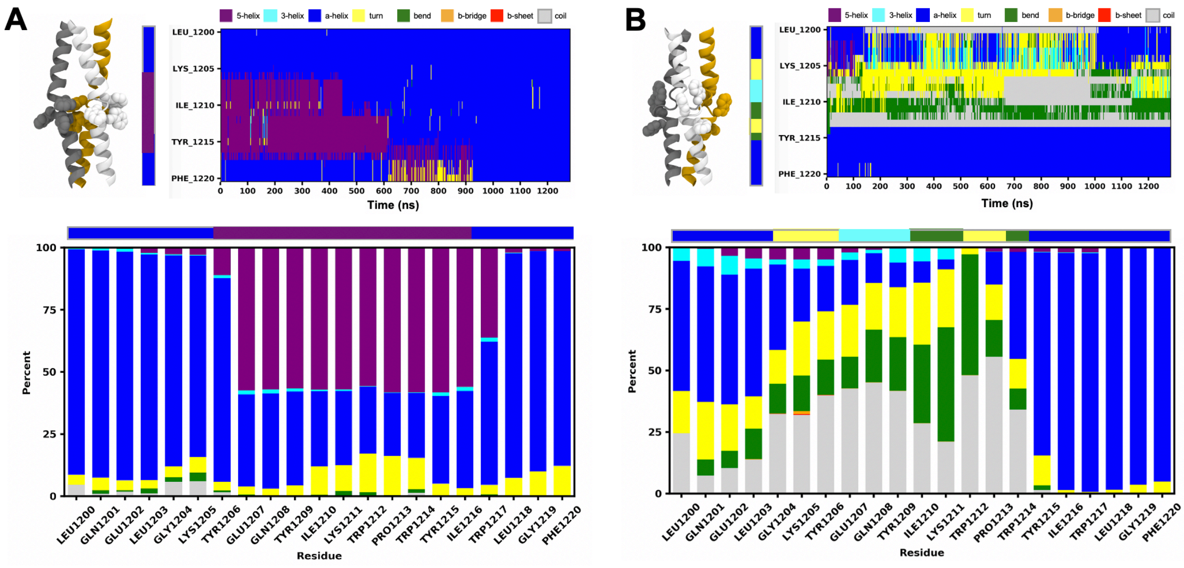


**Figure S3. Secondary structure analysis of HR2-TM linker (A) model 1 and (B) model 2.** (Top from right) a snapshot of model structure, an initial secondary structure, and time series of secondary structures of a representative system, and (bottom) secondary structure contents per residue from all simulations.

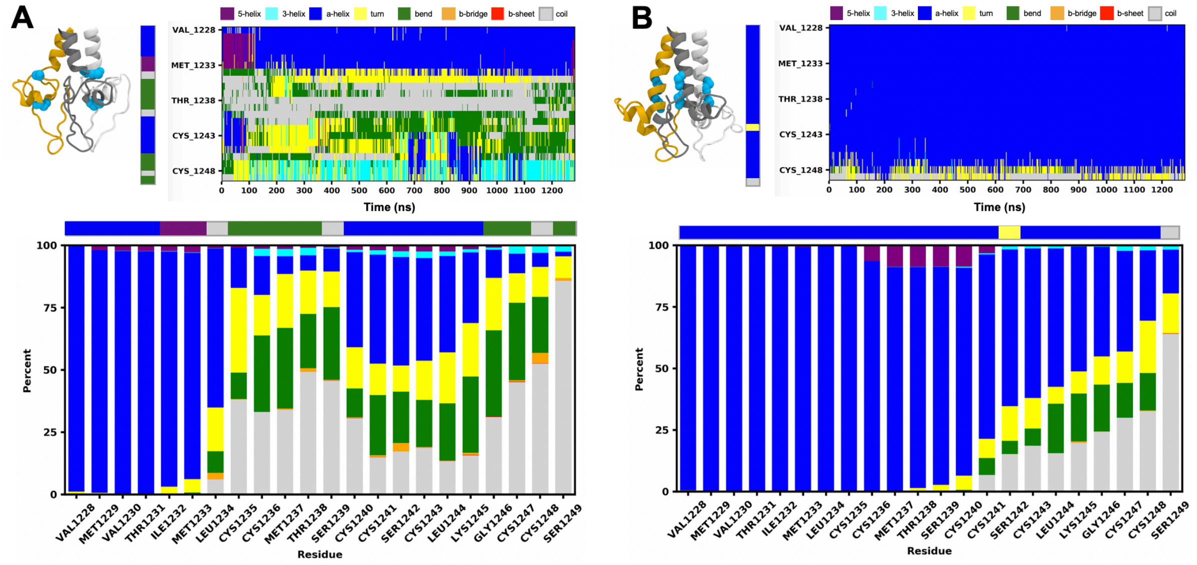


**Figure S4. Secondary structure analysis of CP domain (A) model 1 and (B) model 2.** (Top from right) a snapshot of model structure, an initial secondary structure, and time series of secondary structures of a representative system, and (bottom) secondary structure contents per residue from all simulations.

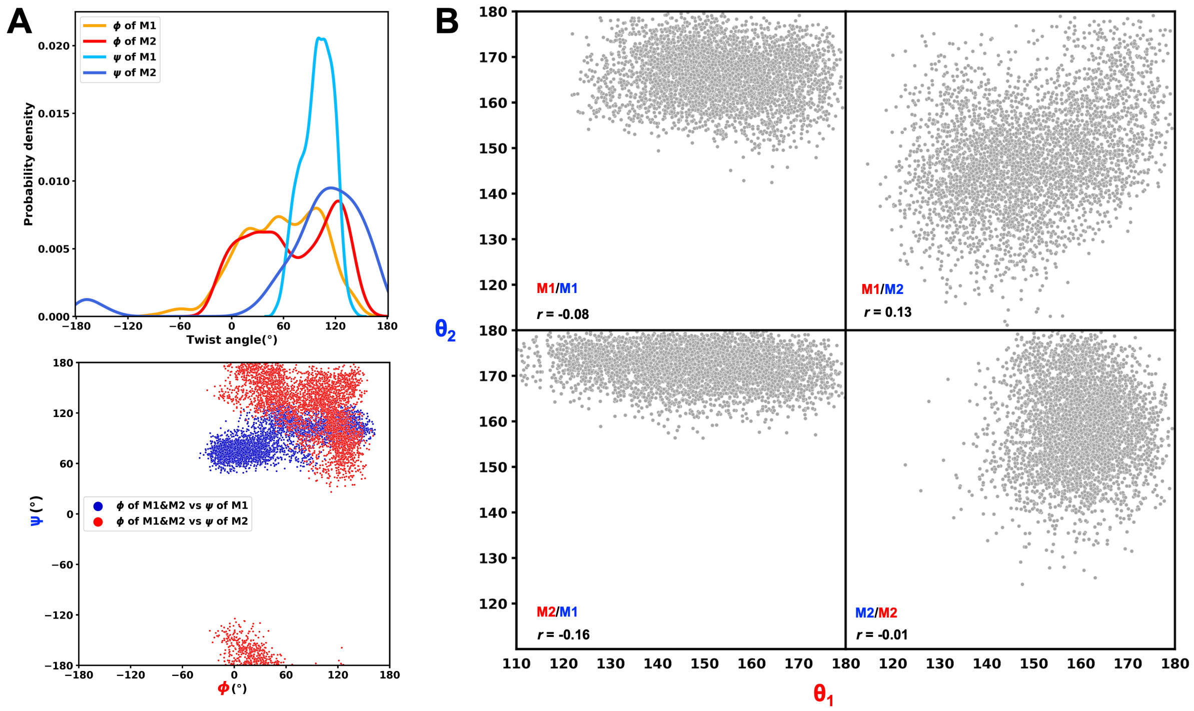


**Figure S5. Twist and bending motions of S protein in a viral membrane.** (A) (top) Probability distribution of twist angle for each HR2 linker and HR2-TM linker model and (bottom) scatter plots of twist angles of HR2 linker and HR2-TM linker. For HR2 linker, both M1 and M2 were used for analysis. (B) Scatter plots of bending angles for HR2 linker and HR2-TM linker models.

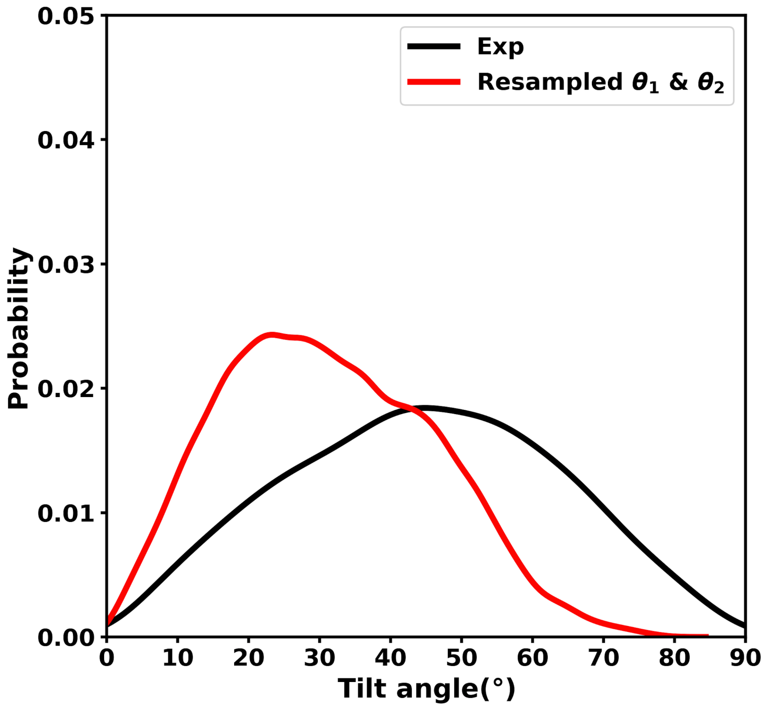


**Figure S6. Tilt angle of S protein in a viral membrane.** Probability distributions of tilt angles for the resampled S protein structures compared to the experimental observation. S protein configurations were resampled based on both M1 and M2 for HR1-HR2 region ($\theta_1$) and only M2 for HR2-TM region ($\theta_2$).

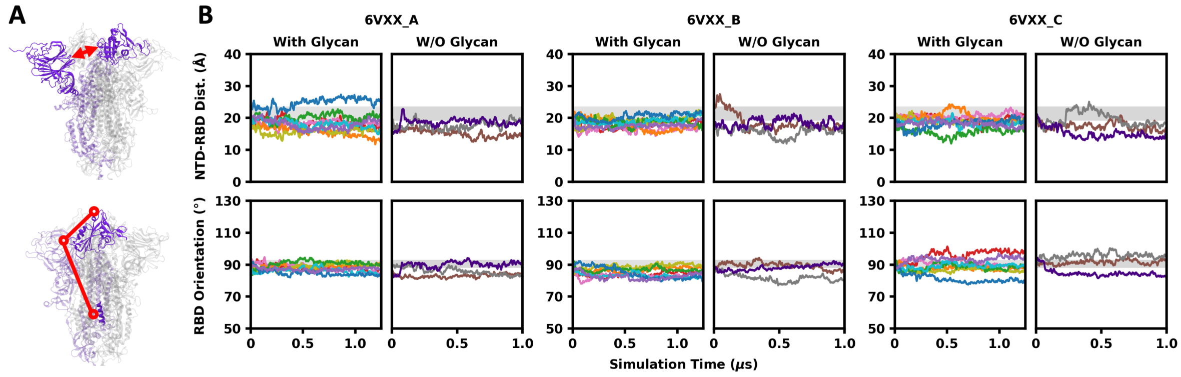


**Figure S7. Motions of RBD and NTD in fully-glycosylated systems and non-glycosylated systems (6VXX).** (A) Illustration of NTD-RBD distance (*d*) and RBD orientation angle (*θ*). (B) The time series of *d* and *θ* in three chains of 6VSB. *d* is defined by the minimum distance between RBD (N334 to P527) and NTD (C15 to S305). *θ* is defined by three points corresponding to the (i) COM of L452 and L492, (ii) COM of N334, and (iii) COM of S1030. The ranges of *d* and *θ* observed in available PDB S protein structures are shaded by gray regions.

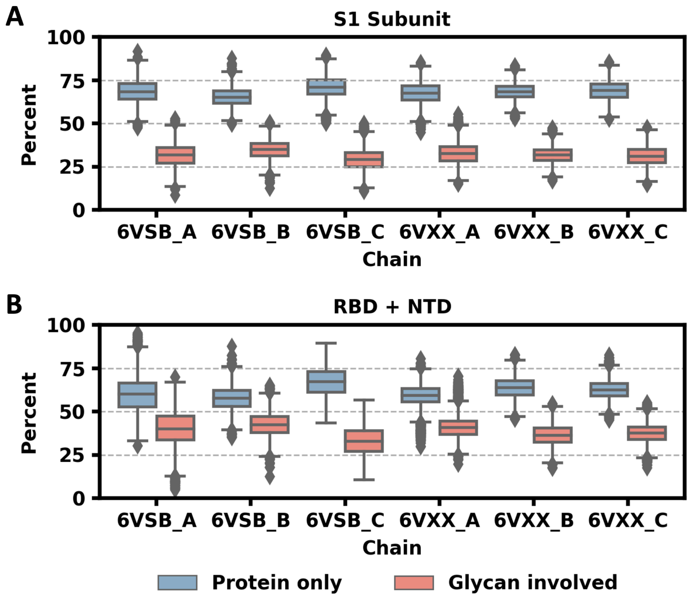


**Figure S8. Surface area buried in the interface of S trimer.** The total buried surface area split into the portion contributed by protein only and the portion involving the contribution from glycans were calculated for (A) the entire S1 subunit and (B) RBD and NTD only.

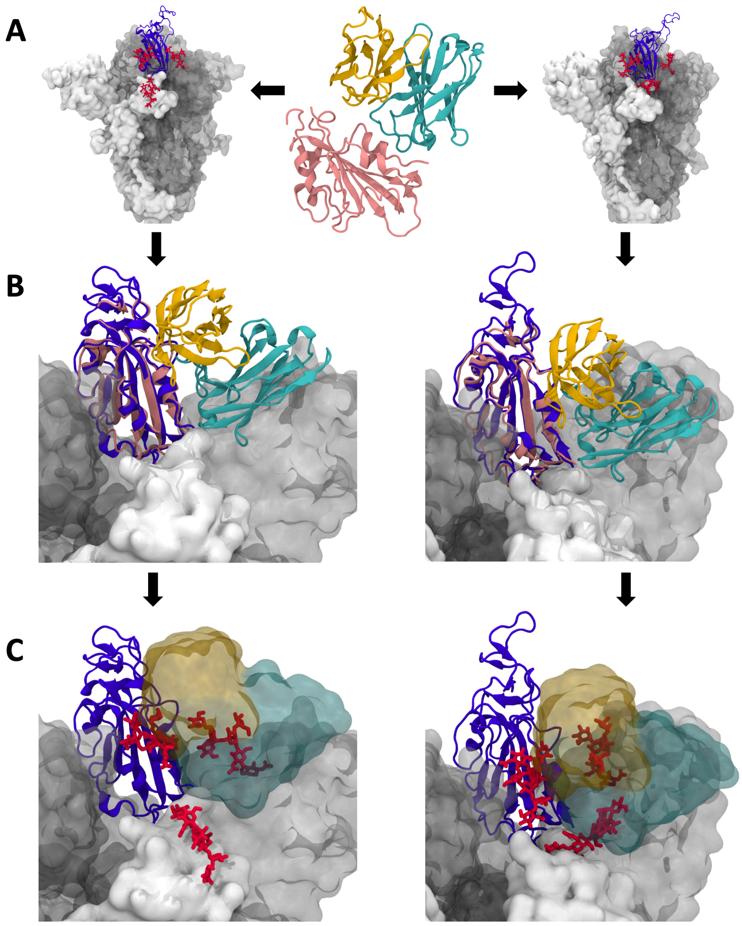


**Figure S9. Procedure of measuring the number of glycan atoms clashing with the superimposed antibody.** (A) Snapshots are extracted from simulation trajectories. (B) The RBD-antibody complex structure is aligned onto the S trimer structure by maximizing the overlap between the RBDs from both structures. (C) The number of glycan heavy atoms clashing with the superimposed antibody (less than 1.0 Å) are counted.

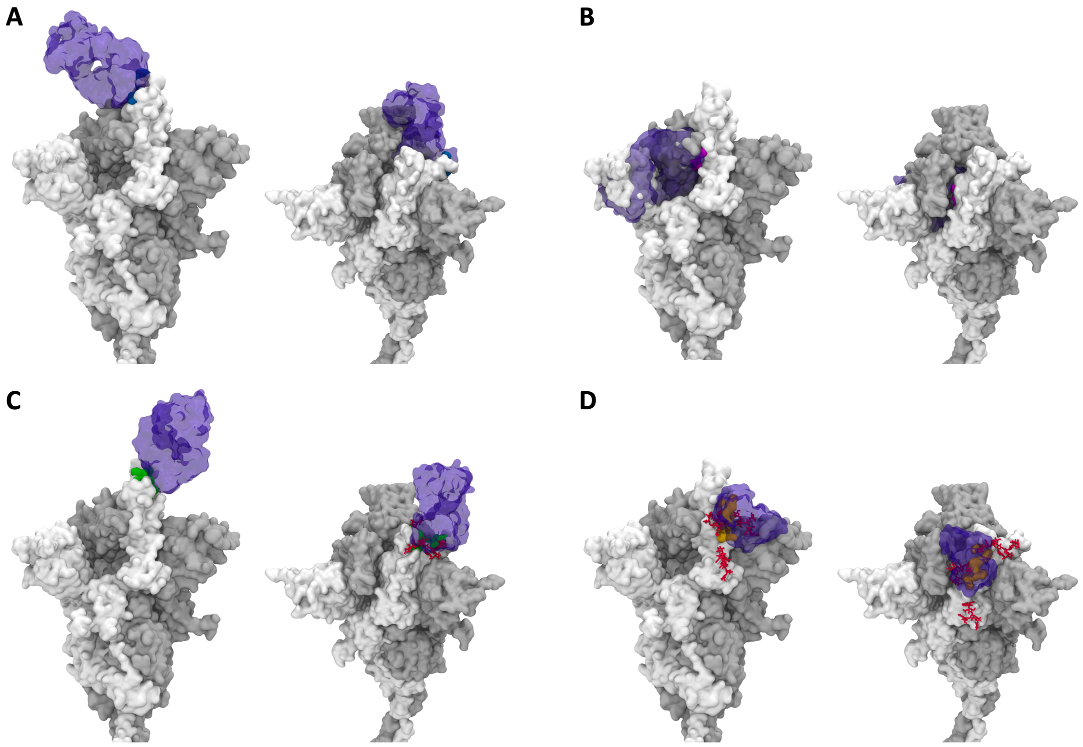


**Figure S10. Four antibodies are aligned onto their epitopes in S trimer.** (A) B38. (B) CR3022. (C) H11-D4. (D) S309. For each antibody, the RBD in the open state is shown in side view (left), and the RBD in the closed state is shown in top view (right).

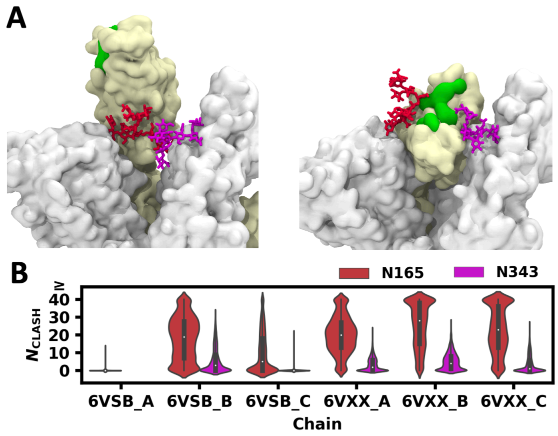


**Figure S11. Clash between glycans and superimposed H11-D4 nanobody.** (A) H11-D4 epitope in RBD (left: open and right: closed), N165 glycan on the neighboring NTD, and N343 glycan on the neighboring RBD. (B) Distributions of glycan heavy atom numbers in clash ($N_{CLASH}$) with the superposed H11-D4 nanobody.

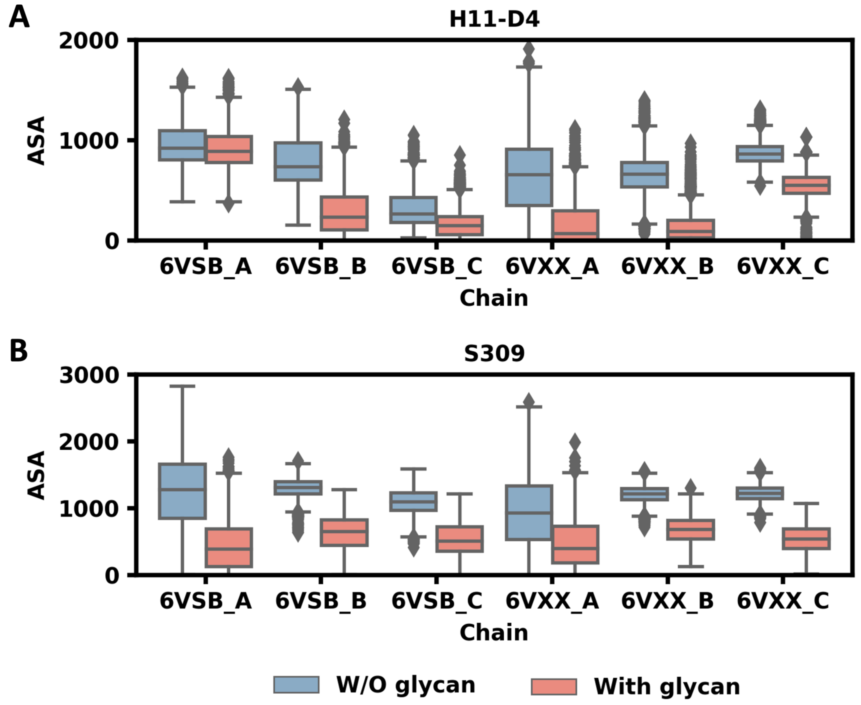


**Figure S12. Accessible surface areas of antibody epitopes when glycans are present or removed.** The ASA of protein portion in the epitopes of (A) H11-D4 and (B) S309 was first calculated with all glycans removed, and then calculated again with all glycans present. In both cases, a probe radius of 7.2 Å was used.
